# Nucleolin acute degradation reveals novel functions in cell cycle progression and division in TNBC

**DOI:** 10.1101/2024.06.17.599429

**Authors:** Joseph Mills, Anna Tessari, Vollter Anastas, Damu Sunil Kumar, Nastaran Samadi Rad, Saranya Lamba, Ilaria Cosentini, Ashley Reers, Zirui Zhu, Wayne O. Miles, Vincenzo Coppola, Emanuele Cocucci, Thomas J. Magliery, Heather Shive, Alexander E. Davies, Lara Rizzotto, Carlo M. Croce, Dario Palmieri

## Abstract

Nucleoli are large nuclear sub-compartments where vital processes, such as ribosome assembly, take place. Technical obstacles still limit our understanding of the biological functions of nucleolar proteins in cell homeostasis and cancer pathogenesis. Since most nucleolar proteins are essential, their abrogation cannot be achieved through conventional approaches. Additionally, the biological activities of many nucleolar proteins are connected to their physiological concentration. Thus, artificial overexpression might not fully recapitulate their endogenous functions.

Proteolysis-based approaches, such as the Auxin Inducible Degron (AID) system paired with CRISPR/Cas9 knock-in gene-editing, have the potential to overcome these limitations, providing unprecedented characterization of the biological activities of endogenous nucleolar proteins.

We applied this system to endogenous nucleolin (NCL), one of the most abundant nucleolar proteins, and characterized the impact of its acute depletion on Triple-Negative Breast Cancer (TNBC) cell behavior.

Abrogation of endogenous NCL reduced proliferation and caused defective cytokinesis, resulting in bi-nucleated tetraploid cells. Bioinformatic analysis of patient data, and quantitative proteomics using our experimental NCL-depleted model, indicated that NCL levels are correlated with the abundance of proteins involved in chromosomal segregation. In conjunction with its effects on sister chromatid dynamics, NCL abrogation enhanced the anti-proliferative effects of chemical inhibitors of mitotic modulators such as the Anaphase Promoting Complex.

In summary, using the AID system in combination with CRISPR/Cas9 for endogenous gene editing, our findings indicate a novel role for NCL in supporting the completion of the cell division in TNBC models, and that its abrogation could enhance the therapeutic activity of mitotic-progression inhibitors.

## Introduction

The nucleolus is a prominent sub-nuclear organelle of eukaryotic cells^1,2^. The most extensively described function of the nucleolus is the synthesis of ribosomal RNA (rRNA) and assembly of the ribosomes^2–4^. The nucleolus also acts as a sequestration compartment for proteins with critical cellular functions such as cell cycle progression, telomere elongation, DNA damage response, and cell death^5^. For these reasons, the nucleolus is now considered a central hub where multiple intra- and extra-cellular signals converge and modulate cell homeostasis and stress response^1,2,5,6^.

Alterations of nucleolar size, structure, and number are associated with physiological aging and a wide range of human diseases, such as neurodegenerative disorders, progeria, and cancer^7–10^. Cancer cells frequently display enlarged nucleoli, possibly to increase ribosome biogenesis to cope with their metabolic needs related to uncontrolled proliferation^3,9,11,12^. In fact, nucleolar parameters, such as increased nucleolar size, are independent prognostic variables in cancer^13^. Particularly in breast cancer, nucleolar area is associated with reduced disease-free survival after surgical tumor resection^13^. For these reasons, nucleolar histologic features are considered an underestimated, clinically relevant indicator associated with poor prognosis^13–15^.

In this study, we investigated the biological activities of nucleolin (NCL), one of the most abundant nucleolar proteins in human cells^16–19^. NCL is involved in rRNA expression and maturation, chromatin remodeling, and translational regulation^16,19–25^. NCL is also consistently upregulated in human tumors, where it has also been found aberrantly expressed on the surface of cancer cells and neoangiogenic blood vessels, but not on their normal/non-proliferative counterparts^16–18,21–23,26,27^. For these reasons, several groups have attempted to develop therapeutic agents targeting NCL on the surface of cancer cells^28–39^. However, deeper understanding of NCL biological functions has remained elusive, which has prevented the clinical translation of anti-NCL agents thus far.

It remains to be determined, for example, whether NCL overexpression has a causal role in cancer pathogenesis or is the result of the transformation process itself. This knowledge gap is partially due to experimental challenges in modulating the expression or the activity of nucleolar proteins in cellular systems. In fact, overexpression of NCL has proven difficult, due to its extreme abundance at the basal level^40^. On the other end, most nucleolar proteins have an essential role, therefore the use of knock-out or shRNA/siRNA approaches, all requiring several days to achieve their effect, is not ideal to assess their direct biological functions^41^. Finally, both overexpression and silencing of nucleolar proteins can lead to a chronic alteration of cell homeostasis, ultimately preventing the ability to discriminate between the biological functions of these proteins and the downstream consequences of their experimental alterations^42^. Overcoming these limitations would improve our understanding of the biological functions of nucleolar proteins, supporting the development of appropriate therapeutic approaches for human diseases where nucleolar functions are altered.

Here, we leveraged the Auxin Inducible Degron *“*on-demand*”* proteolytic system to push the boundaries imposed by NCL knock-down, knock-out, or overexpression approaches^4143–47^. Our *in silico* analyses showed that NCL overexpression is significantly higher in Triple Negative Breast Cancer (TNBC) compared with other breast cancers (BC), which prompted us to use TNBC cell lines for the implementation of this system. Approximatively 15-20% of BC patients are affected by TNBC, a very aggressive subgroup of breast cancer (BC), lacking HER2, Estrogen Receptor (ER) and Progesterone Receptor (PgR), for which no specific treatment is currently available^48–51^. For these patients, the overall survival rate is approximately 77%, but it dramatically drops to 15% for metastatic patients^49^. Every year, approximately 150,000 women die from TNBC, accounting for 30% of all BC-related deaths^52,53^. For these reasons, the identification of molecular determinants of TNBC and of potential therapeutic approaches represents a crucial unmet clinical need.

Using CRISPR/Cas9-based gene editing, we generated TNBC cell lines where endogenous NCL is fused to an Auxin Inducible Degron (AID) tag, without altering its basal expression level. Then, we stably transduced them with the gene encoding for the *Oryza sativa* TIR1 (OsTIR1) E3-ubiquitin ligase protein, which can induce prompt proteasome-mediated degradation of AID-tagged proteins in response to derivatives of the phytohormone Auxin. Given the fast kinetics of this approach (∼ 1 h versus several days using RNAi-based approaches)^43–47,54–57^, the AID system allows the study of protein biological functions with unprecedented specificity and temporal resolution. Then, we performed short-term functional assays to assess the biological impact of NCL acute depletion on the biology of TNBC cells.

Overall, we demonstrate that the acute NCL depletion in TNBC cell resulted in significant changes of the transcriptome and the proteome, altering the cell capability to successfully undergo cytokinesis. These findings support a novel role for NCL in the control of the cell division cycle in TNBC cells.

## Results

### NCL is upregulated in aggressive breast cancer and TNBC

NCL has been previously reported to be upregulated in most human tumors, particularly BC^21,28,38,58^. However, a comprehensive analysis of NCL RNA and protein expression in different subtypes and stages of BC had not been performed. This analysis would facilitate the identification of the most appropriate cancer model to characterize NCL biological functions. We examined patient data collected by The Cancer Genome Atlas (TCGA)^59^, where breast cancer samples are subdivided into the following subtypes based on their gene expression profiling: Normal-like, Luminal A, Luminal B, HER2-enriched, and Basal-Like^59–61^. Our analysis revealed that *NCL* expression levels were significantly higher in the Basal-Like BC subtype, in comparison with Normal-like samples, than any other subtype (**Figure 1A**). A similar analysis, performed on protein data of the CPTAC database^62^, confirmed that Basal-Like BC subtypes expressed the highest levels of NCL protein, in comparison with the others (**Figure S1A**). We also observed a significant increase in NCL protein levels in Basal-Like BC samples in comparison with all the non-basal BC subtypes pooled together (**Figure S1B**).

**Figure 1.**
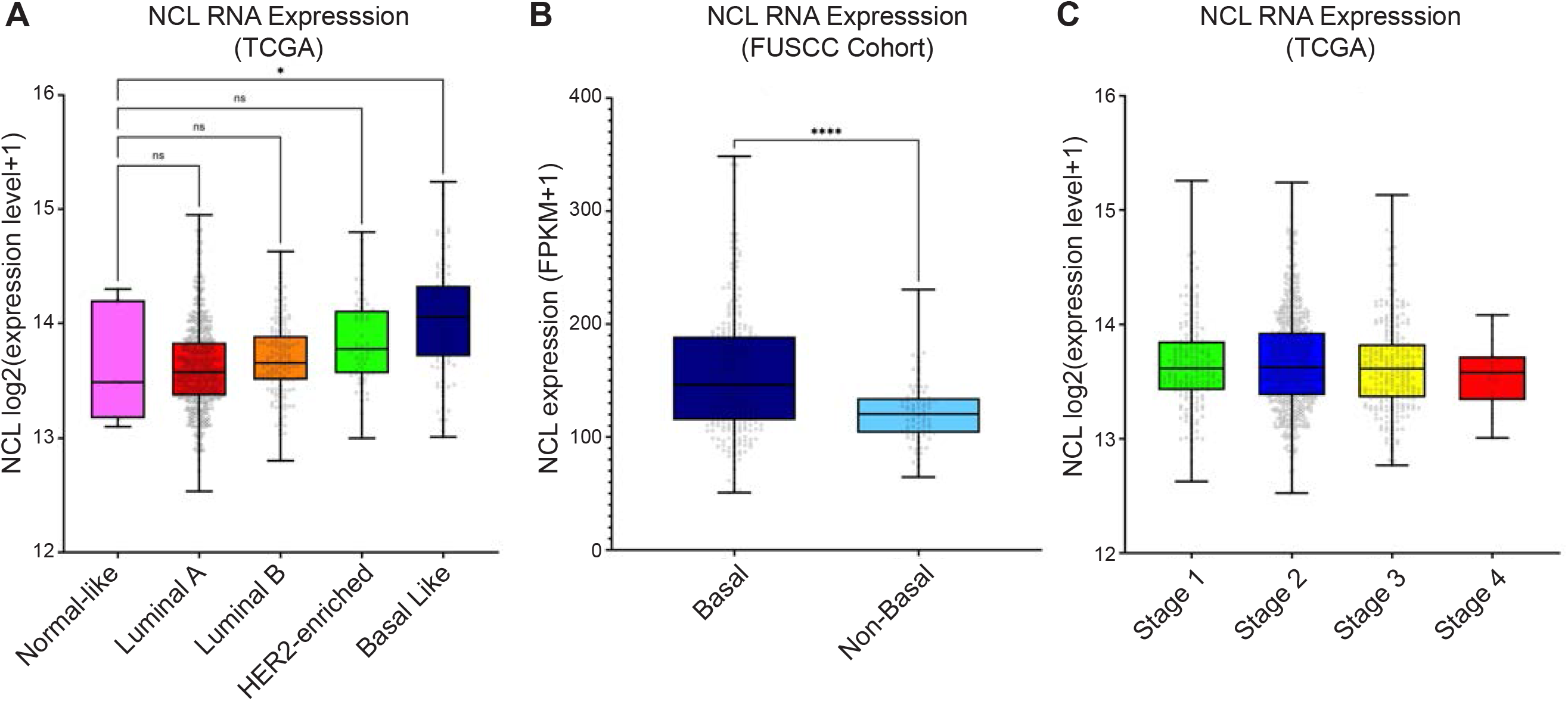
NCL is overexpressed in aggressive human breast tumors. **A)** NCL RNA expression levels among different breast tumor subtypes compared with normal-like breast cancer samples. Data expression levels were obtained from 915 patient samples (Normal-like, n=8; Luminal A, n=624; Luminal B, n=127; HER2-enriched, n=58; Basal Like, n=98) available through The Cancer Genome Atlas (TCGA). Significance was defined using Kruskal-Wallis test. **B)** NCL RNA expression levels in Basal *versus* non-Basal TNBC patients. Data expression levels were obtained from 360 primary Chinese TNBCs (Basal, n=277; Non-Basal, n=83) available through the Fudan University Shanghai Cancer Center (FUSCC). Significance was defined using Mann-Whitney test. **C)** NCL RNA expression levels among 1,076 breast tumor clinical stages, as described in A (Stage 1, n=181; Stage 2, n=624; Stage 3, n=251; Stage4, n=20). Significance was defined using Kruskal-Wallis test. Ns: not significant; *: *p* < 0.05; ****: *p* < 0.0001.

The Basal-Like subtype of BC patients includes the vast majority of TNBC samples^63–66^. Therefore, we assessed *NCL* expression levels on an independent dataset (FUSCC cohort^66^) of TNBC patients, previously stratified as Basal (characterized by an aggressive phenotype and poor prognosis) or non-Basal (less aggressive and with a better prognosis). Our analysis indicated that NCL levels were significantly higher in the Basal subtype of TNBC (**Figure 1B**). Finally, we queried the TCGA database to assess whether *NCL* overexpression was associated with specific clinical stages of BC development. As shown in **Figure 1C**, BC samples classified as Stage I through IV did not display a significant association between NCL overexpression and BC clinical stage.

Altogether, these results indicate that, among all BCs, Basal-Like BCs, and specifically TNBC with worse prognosis and aggressive phenotype, display higher NCL RNA and protein levels. Notably, NCL expression did not change during clinical progression, suggesting that its overexpression might be associated with the early stages of BC pathogenesis or initial oncogenic events.

### The Auxin Inducible Degron system accomplishes an efficient and fast depletion of NCL

Since NCL is significantly upregulated in TNBC, we decided to investigate the effects of its complete abrogation in a cellular model of this BC subtype. We used the Cancer Dependency Map (DepMap)^67,68^ portal to identify the most suitable TNBC cell line for our study. This score measures the effect of knocking out a gene on cell viability and is derived from large-scale CRISPR or RNAi screens across numerous cancer cell lines ^53,54^. DepMap defines genes as commonly essential when their perturbation score is in the top 10% of all genes, and a dependency score of −0.5 or lower is considered a significant dependency. As shown in **Figure S2A-B,** NCL was considered a commonly essential gene both using RNAi-based systems (Achilles+DRIVE+Marcotte, and DEMETER2 databases, 487/710 cell lines) and CRISPR-based knock-out (DepMap Public 23Q4+ Score, and Chronos databases, 1123/1150 cell lines), not only in BC cell lines, but in any available cell line, with no cell line with a CRISPR-KO perturbation score higher than −0.16 (LB771HNC Head and Neck Squamous Cell Carcinoma cell line). For breast cancer cell lines, the highest observed score was −0.52 (MDA-MB-361 cells, an ER-positive/PgR-negative luminal mammary carcinoma cell line^69^). Notably, a lower overall score was observed when using siRNA in comparison with CRISPR-based KO, probably due to differences in silencing efficacy between the two approaches^70^. These observations strongly suggest that the generation of a cancer cellular system where NCL is completely and chronically abrogated is either not possible or would require significant adaptation phenomena for the cells to survive. Therefore, results obtained from chronic knockdown experiments could be biased by artifacts due to cellular compensatory processes not directly related to NCL abrogation.

We reasoned that a targeted proteolysis system would be the best tool to achieve a rapid, “on-demand” degradation of NCL, and we decided to take advantage of the Auxin Inducible Degron (AID) system^43–47,54^ in combination with CRISPR/Cas9-based knock-in gene editing to modify the endogenous *NCL* gene.

The AID system has never been used for endogenous proteins with nucleolar localization to date, but it could allow the study NCL biological functions with unprecedented temporal resolution. As a cellular model, we chose MDA-MB-231, a widely used TNBC cell line, expressing high levels of endogenous NCL^30,38^. This specific cell line was also selected based on previous studies investigating the impact of NCL inhibition in breast cancer cell lines^71–73^. Moreover, the genomic location of the human *NCL* gene (chr2, q37.1) is not involved in chromosomal alterations in MDA-MB-231 cells^74,75^, suggesting that this cell type has only 2 copies of our gene of interest. The absence of copy number variations (CNVs) of the *NCL* in this cell line was also confirmed by the data available in the DepMap portal (**Figure S2C**)

We employed a two step-editing CRISPR/Cas9-based knock-in approach to generate cells where endogenous NCL protein could be quickly degraded on demand. For the targeted genomic modification of the endogenous NCL, we engineered a plasmid containing two 1-Kb homology arms corresponding to the genomic region of the 5’-end of the endogenous *NCL* coding sequence (**Figure 2A**). The choice of the 5’-end of the *NCL* gene was based on previous reports indicating that modifications of NCL N-terminal region does not affect its biological functions^76^. We included in this plasmid the mAID2 tag^44,45,47,54^, in frame with either the mCherry2^77^ or the HaloTag^78^ proteins. Both these two plasmids were used as donors for Cas9-based homology-driven repair (HDR) of the endogenous *NCL* gene. We decided to use two different constructs for gene-editing purposes because this strategy offers multiple advantages: first, it leads to the expression of endogenous NCL in frame either with the two fluorescent modules (mCherry2 and HaloTag), allowing the tracking of its expression and localization; second, it allows for the selection of cells displaying a multi-allelic *NCL* editing, based on the presence of both fluorescent tags (**Figure 2B**). Following successful targeting, we then transduced the homozygous AID-NCL cells with a construct containing the *OsTIR1*, constitutively integrated in the *AAVS1* safe-harbor locus^79^. Specifically, we generated transgenic cell lines expressing OsTIR1(F74G) (a.k.a. OsTIR1v2), a point-mutant variant of the *Oryza Sativa* TIR1, whose activity is regulated by the auxin synthetic derivative molecule 5-phenyl-Auxin (5-Ph-IAA)^47^. Compared to its initial version, the OsTIR1(F74G) displays reduced leakiness, and it is more sensitive (500-fold) to its synthetic ligand ^47^.

**Figure 2.**
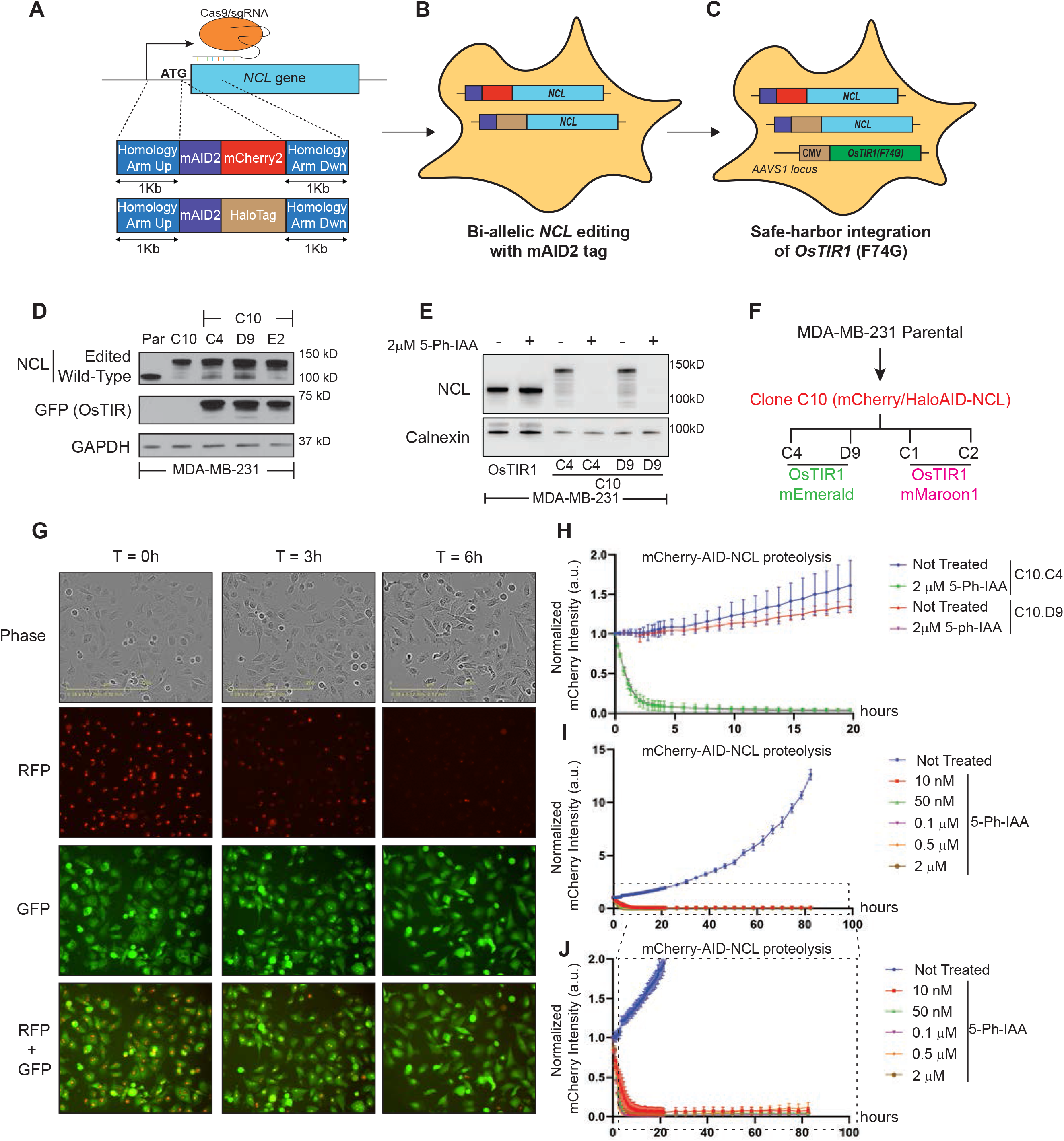
The Auxin Inducible Degron system accomplishes an efficient and fast depletion of NCL. **A-C)** Schematic representation of the approach used for the generation of homozygous AID-NCL/OsTIR1 cell lines. Multi-allelic editing of endogenous *NCL* gene was achieved using two independent donor plasmids and a gRNA designed against the first translated codon of the gene (**A**). Selected multi-allelic *NCL* edited cells (**B**) were further modified to integrate the *OsTIR1* transgene into the *AAVS1*-safe harbor site (**C**). See also Figure S2. **D**). Western blot of endogenous NCL in engineered clones, in comparison with parental (Par) cells. Anti-GFP antibody was used to detect the OsTIR1 fused to mEmerald. GAPDH was used as loading control. **E)** NCL degradation by western blot at 24 h upon 5-Ph-IAA treatment in two different MDA-MB-231 clones (C10.C4 and C10.D9) compared with MDA-MB-231 cells only transduced with OsTIR1. Calnexin was used as loading control. **F)** Diagram of the hierarchical generation of the clones used in the current study. **G)** Representative images of Incucyte experiments on MDA-MB-231 mCherry2/Halo-AID-NCL/OsTIR1-mEmerald cells. The RFP channel was used to detect mCherry2-AID-NCL, and the GFP channel was used to assess OsTIR1-mEmerald expression and localization. **H)** Quantitative analysis of mCherry (AID-NCL) intensity from Incucyte experiments on the indicated cell clones. RFP integrated intensity per image was normalized for t=42’ to account for potential plate condensation at the early time points. The experiment was performed in 5 biological replicates, with 5 technical replicates. Error bars show standard deviation. **I-J**) Dose-response curve (**I**) and relative magnification (**J**) for mCherry-AID-NCL degradation using the indicated amounts of 5-Ph-IAA. RFP integrated intensity per image was normalized for t=0. The experiment was performed in three biological replicates, with four technical replicates. Error bars show standard deviation.

We generated cells expressing OsTIR1(F74G) fused to the bright green fluorescent protein mEmerald (a variant of GFP, with enhanced brightness^80^). (**Figure 2C and S2D**), to simplify the identification of transduced cells. Importantly, this chimeric protein allows to confirm the expression and the localization of OsTIR1(F74G). Western blot analyses confirmed that gene-edited MDA-MB-231 (Clone C10) expressed the AID-NCL fusions, but not the wild-type counterpart. Moreover, C10 subclones (C10.C4, C10.D9, and C10.E2) also expressed OsTIR1v2 (detected using an anti-GPF/mEmerald antibody) (**Figure 2D**). As expected, treatment with 2 μM 5-Ph-IAA for 24 h completely abrogated NCL to almost undetectable levels in both clones (**Figure 2E**). This result also allowed us to confirm that the lower molecular weight bands detected by WB analyses correspond to AID-tagged NCL, potentially due to proteolysis or presence of post-translational modifications^19^, as their presence was completely abrogated upon 5-Ph-IAA treatment.

To generate a novel AID system, compatible with other green channel-based experimental techniques, we generated an integration construct which co-expresses OsTIR1(F74G) and a histone H1.0 fluorescently tagged with the far-red protein mMaroon1^81^ (**Figure S2E**). This alternative version allows a simultaneous visualization of chromatin/nuclei, and their shape grants the discrimination of cells undergoing M phase^81^.

We obtained two additional independent clones (C10.C1 and C10.C2, summarized in **Figure 2F**), where we confirmed OsTIR1 expression (**Figure S2F)** and acute NCL degradation (**Figure S2G**). NCL abrogation was also observed by live-cell imaging experiments (**Figure 2G** and **S2H**). We quantified the total integrated RFP signal (corresponding to the AID-mCherry2-NCL allele) over time up to 20 h, confirming NCL abrogation at approximately 6 h of 5-Ph-IAA treatment in all the analyzed clones (**Figure 2H and S2I**). We also observed that OsTIR-mediated AID-mCherry2-NCL degradation was only minimally dependent on the 5-Ph-IAA dose (**Figure 2I**, and magnification in **Figure 2J**), and persisted up to 80 h after a single treatment. Live-cell imaging of OsTIR1-mEmerald (GFP channel) revealed enhanced localization (0-3 h) of the enzyme at the nucleoli following 5-Ph-IAA treatment (**Figure 2G** and **S2H**). However, in untreated cells, the enzyme had a homogeneous distribution over the cell volume, as previously described^82^.

Altogether, these data demonstrate the effectiveness of the AID system in quickly degrading endogenously edited NCL.

### The acute abrogation of NCL reduces cell proliferation and alters the RNA levels of genes involved in cell cycle progression

Previous reports have shown that NCL silencing results in cell cycle progression defects. However, these studies have shown contrasting results with accumulation of cells either in G1 or G2/M phase of the cell cycle^41,83^. We hypothesize that this discrepancy could be due, at least in part, to the limited efficacy and temporal resolution of the silencing system used in these studies. Therefore, we decided to test whether acute abrogation using our AID system could elucidate the role of NCL in cell cycle progression.

To validate the impact of NCL abrogation on cell proliferation, we performed cell growth assays using Incucyte live-cell imaging in the presence or in the absence of 5-Ph-IAA to induce acute NCL depletion. NCL abrogation resulted in a marked reduction in proliferation, but only at 48 h from the treatment (**Figure 3A**, **Figure S3)**. Conversely, all control conditions (**Figure 3B, Figure S3**) confirmed that neither 5-Ph-IAA treatment, nor NCL editing with the AID tag, nor OsTIR1 expression had any statistically significant effect on cell proliferation when all the components of the AID system were available.

**Figure 3.**
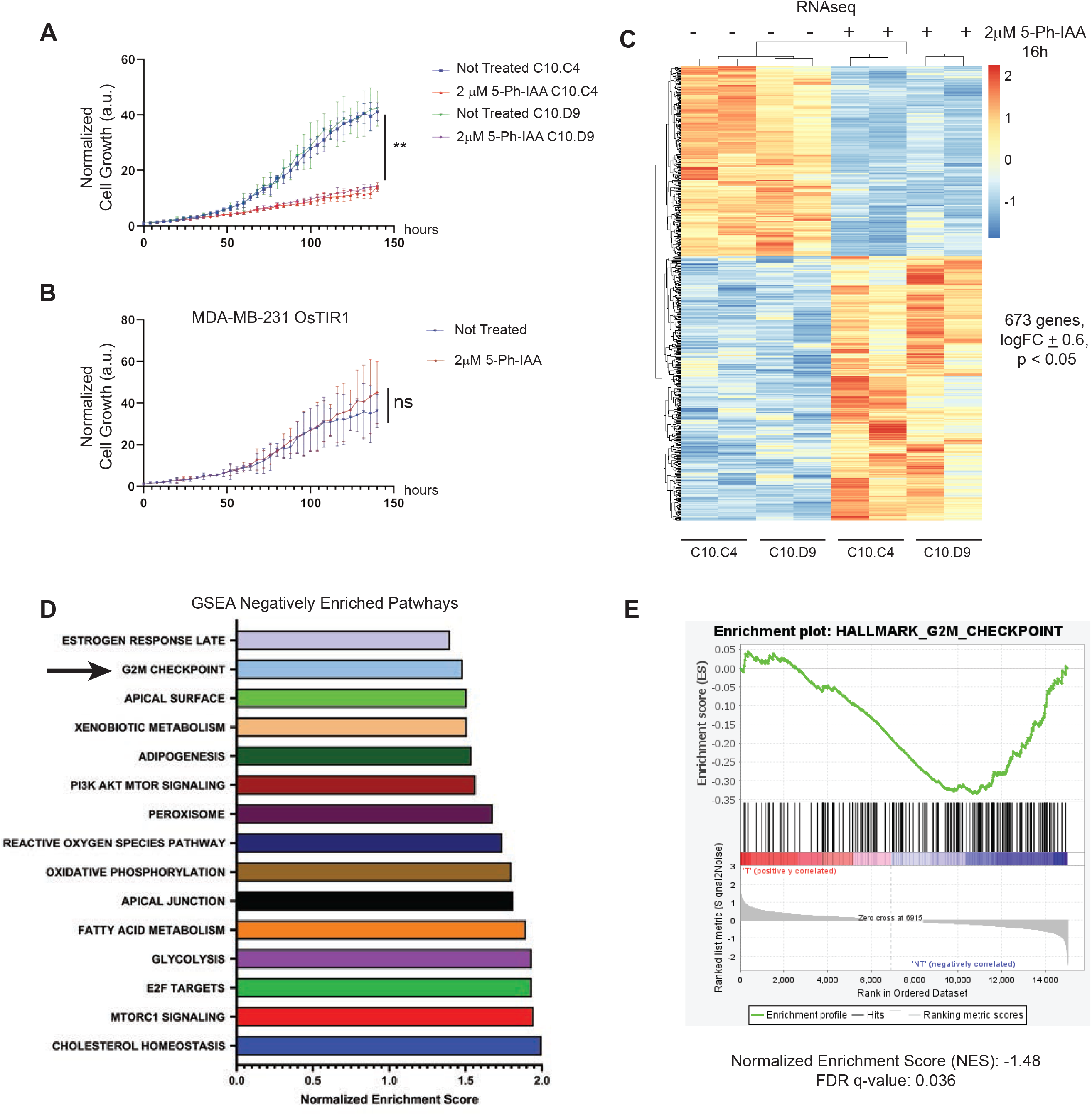
NCL acute abrogation alters the gene expression profile and proliferation of TNBC cells. **(A-B)** Growth curve of MDA-MB-231 OsTIR1/AID-NCL or control MDA-MB-231 OsTIR1 clones treated with 2μM 5-Ph-IAA or left untreated for up to 140 hours. Incucyte live-cell imaging system was used to monitor cell proliferation. The experiment was performed in five biological replicates, with 5 technical replicates. Statistical significance was calculated using 2-way ANOVA with Tukey’s multiple comparison test. Error bars show standard deviation. **: *p* < 0.01; ns: not significant. **C)** Heatmap of RNA sequencing analyses on NCL-edited clones at 16 h upon 2μM 5-Ph-IAA treatment or left untreated. Two independent clones in biological duplicate were used for the experiment. **D)** Gene set enrichment analysis (GSEA) of Hallmark Pathways using RNA sequencing data of 5-Ph-IAA treated versus non-treated NCL-degron clones. Ranking of Normalized Enrichment Scores of the 15 most negatively enriched pathways after NCL abrogation. **E)** Enrichment plot for the G2/M Checkpoint pathway, as described in D.

NCL is an RNA-binding protein, with the capability of modulating RNA expression, stability, and translation^16–19^. Therefore, we decided to investigate whether the effect of NCL abrogation on cell proliferation could be due to alteration of RNA abundance, via RNA-sequencing. To this aim, we collected and analyzed total RNA from MDA-MB-231 AID-NCL cells treated with 2 μM 5-Ph-IAA for 16 h. This time point was selected to maximize NCL degradation while minimizing potential indirect effects on RNA abundance. As shown in **Figure 3C**, 673 genes were significantly de-regulated upon NCL abrogation, including 390 genes upregulated and 283 genes downregulated (p < 0.05, LogFC ±0.6). This relatively low number of genes is consistent with the short-term duration of this experiment, and potentially includes genes that are immediately associated with a direct regulation by NCL. Gene Set Enrichment Analysis (GSEA) identified several known genes whose expression is regulated by NCL, such as E2F targets and members of the AKT/mTOR signaling^84,85^ (**Figure 3D**). Among the most de-regulated pathways (including MTORC1 signaling and PI3K/AKT/MTOR signaling), we identified genes involved in G2/M cell cycle checkpoint (**Figure 3E**, Normalized Enrichment Score: −1.48; FDR q value: 0.036). Our experimental approach demonstrated that this could be related to immediate changes in the abundance of specific genes associated with G2/M transition.

These experiments demonstrated for the first time that acute NCL depletion has a rapid and negative effect on the expression of genes regulating cell proliferation.

### NCL abrogation causes cytokinesis defects

Next, we decided to determine which phase of the cell cycle was most impacted by NCL acute abrogation, ultimately resulting in reduced cell proliferation. To this end, we treated MDA-MB-231 AID-NCL/OsTIR1/H1.0-mMaroon1 cells with 5-Ph-IAA, which were collected at different time points, and analyzed for their content in DNA and phospho-histone H3(S10), a known marker of active mitosis, to differentiate cells in G2 or M phase^86^.

Propidium iodide staining showed a significant increase in cells with a tetraploid (4N) amount of DNA at 48 and 72 h after NCL abrogation (**Figure 4A-B**). The 4N population was identified by our ModFit cell cycle deconvolution as a potential G2/M arrest. However, we did not see a proportional increase in phospho-Histone H3(S10) (**Figure 4C-D**), but rather a decrease in this population, indicating that NCL abrogation does not result in a prolonged arrest in the active phase of mitosis. These results are consistent with two possibilities: either NCL-depleted cells would accumulate in the G2 phase, at the end of the complete replication of DNA (S/G2 transition phase) but before the transition into M phase; or, in the absence of NCL, cells could accumulate in G1, but with a tetraploid content of DNA. To distinguish between these alternative hypotheses, we performed immunofluorescence staining of LaminA/C (a marker of nuclear lamina^87^), and actin filaments (using Alexa-647 Phalloidin) to identify cell boundaries, after 72 h of 5-Ph-IAA treatment. NCL abrogation caused a significant increase in bi-nucleated cells in both clones, but not in control cells containing OsTIR1 and wild-type NCL, upon 5-Ph-IAA treatment (**Figure 4E-F**).

**Figure 4.**
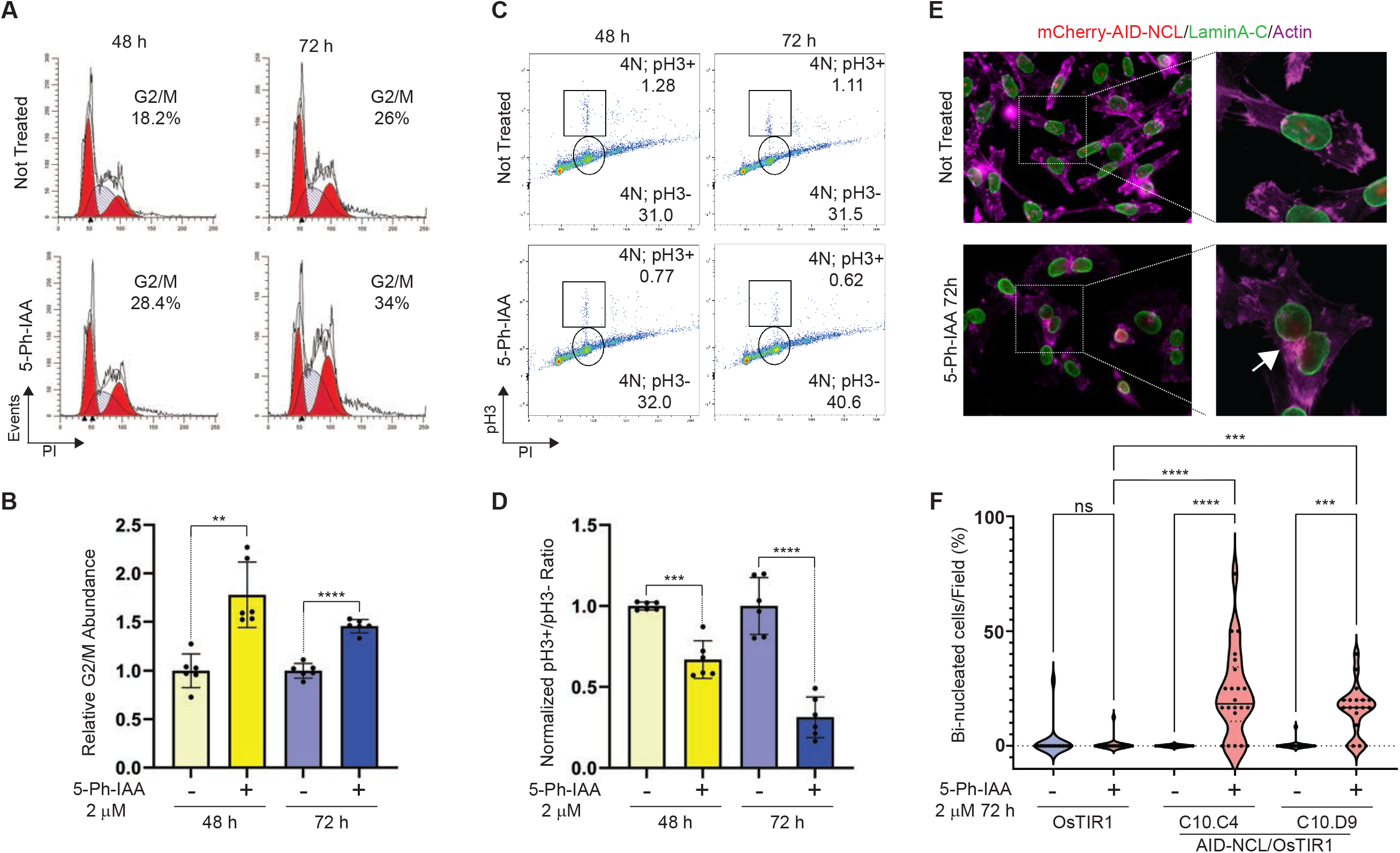
NCL acute abrogation induces defects of cell cycle progression and cytokinesis. **(A)** Representative flow cytometry data of cell cycle analyses by propidium iodide staining of OsTIR1/AID-NCL-edited MDA-MB-231 clones. The percentage of cells in each phase of the cell cycle was obtained by curve deconvolution using ModFit. The absolute abundance of cells in G2/M phase for the specific representative images is also reported. **(B)** Quantitative analyses of the relative percentage of cells in G2/M phase, as shown in A. Data are representative of two independent clones analyzed in three experiments performed in technical duplicate and normalized for the percentage of cells in G2/M in untreated controls. Significance was calculated using Welch’s t-test. **(C)** Representative flow cytometry data of propidium iodide/phospho-Histone H3 (S10) staining in cells. Gating was designed to quantitate the absolute population of cells with a 4N content of DNA, and either positively (4N; pH3+) or negatively (4N; pH3-) stained for pH3(S10) antibody. **D)** Quantitative analyses of the 4N; pH3+/4N; pH3-abundance ratio, as shown in C. Data are representative of two independent clones analyzed in three experiments performed in technical duplicate and normalized for the untreated controls. Significance was calculated using Welch’s t-test. **(E)** Representative images of immunofluorescence images of AID-NCL edited MDA-MB-231 clones after the treatment with 2μM 5-Ph-IAA for 72 h or left untreated. LaminA/C staining (green) was used to detect nuclear membrane. Phalloidin-iFluor 647 staining (magenta) was used to stain actin cytoskeleton membrane. Edited mCherry2-AID-NCL is also visible in untreated cells (red). White arrow indicates actin patches in a bi-nucleated cell. **(F)** Quantification of bi-nucleated cells identified in the experiments shown in E. For each sample, at least 15 images, representative of independent fields, were taken on a single focal plane at 60x magnification. A total of at least 100 cells were captured for each sample. The percentage of bi-nucleated cells per image was used to generate the violin plots. Statistical significance was calculated using one-way ANOVA with Sidak’s multiple comparison test. Data are representative of one of two independent experiments. **: *p* < 0.01; ***: *p* < 0.001; ****: *p* < 0.0001; ns: not significant.

Collectively, these results show that the acute abrogation of NCL impairs cell cycle progression and causes an accumulation of tetraploid bi-nucleated cells, suggesting that NCL could play a role in preventing cytokinesis failure^71^.

### NCL expression is associated with proteins involved in chromosome segregation

So far, we have demonstrated that the acute depletion of NCL causes defects of cell proliferation and alterations of the transcriptome, pointing to a possible involvement of NCL in cytokinesis. Defects of physical cell separation of the two daughter cells at the end of mitosis are defined as cytokinesis failure (reviewed in^88^). The best characterized causes of cytokinesis failure are: 1) defects/mutations of cytokinesis regulators; 2) long delays in mitosis (mitotic slippage); and 3) physical obstructions preventing the furrow cleavage after mitosis^88^. The transcriptome from TNBC NCL-depleted cells pointed toward mitotic slippage as the probable cause of cytokinesis failure, due to the reduced expression of genes involved in G2/M transition (**Figure 2A-C**). However, delayed mitosis usually results in the formation of cells with a single tetraploid nucleus, or multiple micronuclei, which we did not observe in our TNBC NCL-depleted cells (**Figure 4**)^88^. Furthermore, binucleated cells after NCL depletion showed the accumulation of actin patches (**Figure 4E, right bottom panel**), which have been previously shown as the result of the formation of chromosomal bridges, lagging chromosomes, and altered chromosomal segregation^89^.

Therefore, we hypothesized that higher NCL levels could be associated with increased levels of proteins involved in chromosomal dynamics in TNBC. To validate this hypothesis, we analyzed the TCGA database (Harvard Firehose Initiative, >1,000 breast invasive carcinomas) to reveal the transcripts positively correlating with NCL expression. The Spearman’s Correlation level for each gene was used to generate an ordered gene rank list, further analyzed by GSEA (**Figure 5A** and **S4A**). As expected, based on the known NCL roles in these processes^16,18,19,21,24^, the most enriched GO terms were “Ribosome Biogenesis” and “Ribonucleoprotein Complex Biogenesis”. However, five out of ten of the most enriched pathways included DNA dynamics processes, such as “Chromosome Segregation” and “Sister Chromatid Segregation”. Encouraged by these results, to assess whether NCL expression positively correlated with genes involved in DNA dynamics, we performed a parallel analysis of publicly available CPTAC proteomics data. All the ten pathways identified in the correlative analysis of the transcriptome were also highly correlating at protein level with the expression of NCL protein (**Figure 5B**).

**Figure 5.**
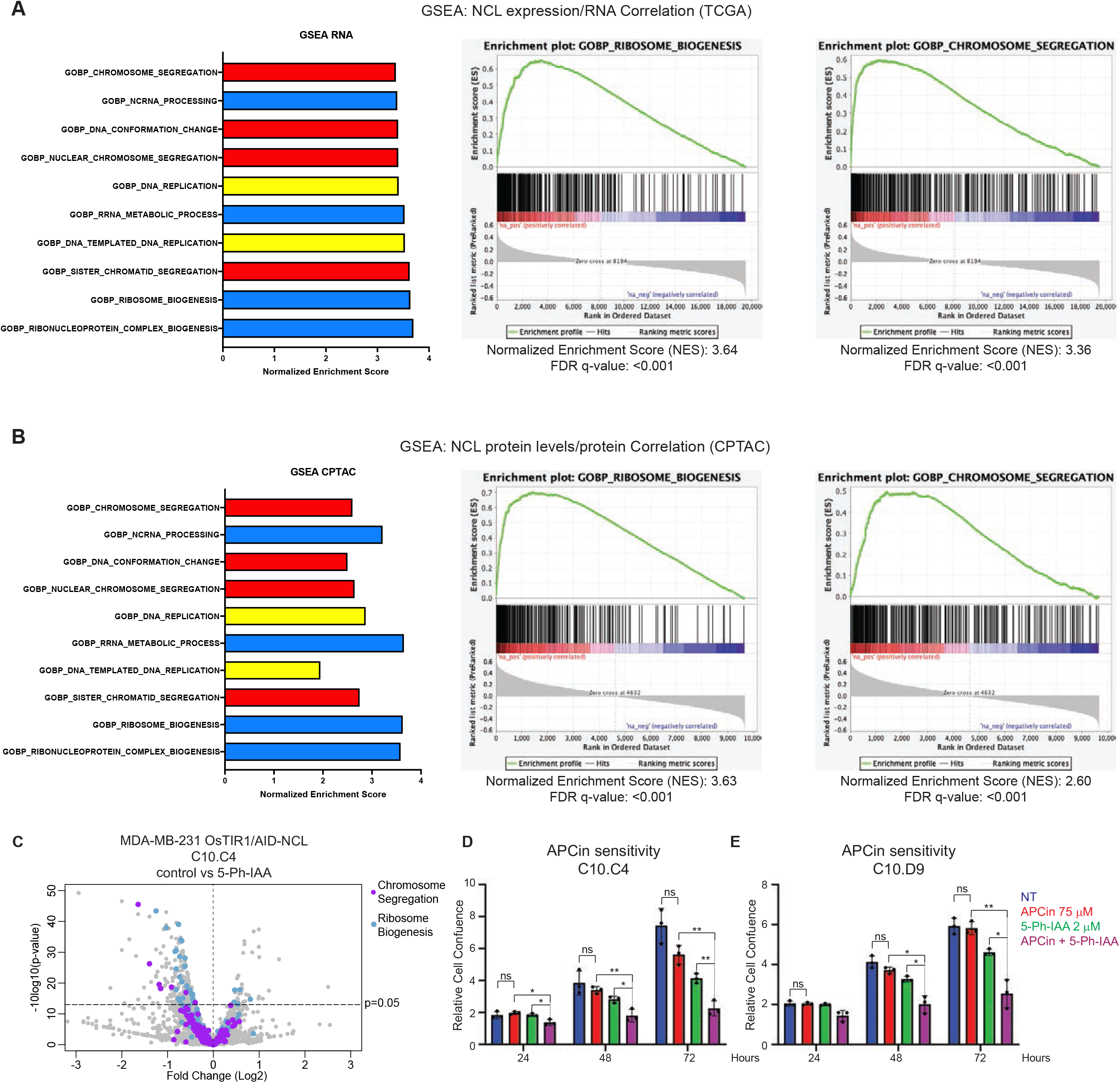
NCL levels are associated with altered sister chromatid segregation and sensitivity to inhibition of the Anaphase Promoting Complex. **A)** Gene set enrichment analysis of GO Terms by gene ranking based on their correlation with NCL mRNA levels in the TCGA database. The top ten most enriched pathways positively correlated with NCL RNA expression are represented. Enrichment plots of two representative GO Terms are also reported (see also Figure S4). **B)** Gene set enrichment analysis of GO Terms identified in A were analyzed based on their correlation with NCL protein levels in the CPTAC database. Enrichment plots of two representative GO Terms (as in A) are also reported. **C)** Volcano plot of differentially abundant proteins upon NCL abrogation in MDA-MB-231 OsTIR1/AID-NCL cells (clone C10.C4), detected by TMT-proteomic analysis. Reported data excludes proteins with −10log_10_(p-value)> 50, for better visualization. For the full range, see Figure S4. Proteins included in the GO Terms “Chromosome Segregation” and “Ribosome Biogenesis” are reported in purple and blue, respectively. **D-E)** Assessment of cell proliferation of two independent MDA-MB-231 OsTIR/AID-NCL clones, left untreated or treated with indicated doses 5-phIAA, APCin, or combination of both, for the indicated time points. Growth was assessed by Incucyte live-cell imaging and reported as relative growth normalized vs t=36’. See also Figure S4. The experiment was performed in three biological replicates, with five technical replicates. Error bars show standard deviation. Statistical significance was calculated using 2-way ANOVA with Sidak’s multiple comparison test. Ns: not significant; *: *p* < 0.05; **: *p* < 0.01.

Then, we decided to assess in our TNBC in vitro model whether the acute depletion of NCL had any negative impact on the abundance of the proteins identified *in silico*. To this aim, we performed a Tandem Mass Tag quantitative proteomics analysis on MDA-MB-231 AID-NCL/OsTIR1 cells at 24 h after 5-Ph-IAA treatment to assess the global protein de-regulation upon NCL acute degradation. As expected, the protein with the most statistically significant downregulation upon 5-Ph-IAA was NCL. Results also showed that there was general downregulation of proteins involved in Ribosome Biogenesis (**Figure S4B, Figure 5C**, blue dots, and **Supplementary Table 1**). Additionally, mimicking the patient correlative analysis (**Figure 5A-B**), seven proteins involved in Chromosome Segregation (**Figure 5C**, purple dots, and **Supplementary Table 2**) were significantly downregulated (p < 0.05) after NCL degradation, while no member of this GO term family was upregulated.

These results confirmed that depletion of NCL had a negative impact on factors involved in chromosomal segregation, potentially explaining the observed defects on cell cycle progression (**Figure 3 and 4**).

Chromosomal segregation is regulated by the activity of the Anaphase Promoting Complex/Cyclosome (APC/C) and its regulatory subunit Cdc20^90,91^. Inhibition of APC/C^Cdc20^ is synthetically lethal with defective sister chromatid cohesion^92^. Therefore, we hypothesized that altered sister chromatid dynamics upon NCL abrogation could enhance the activity of APC/C^Cdc20^ inhibitors, such as APCin^93^. To test this, we performed growth curves on MDA-MB-231 NCL-AID/OsTIR1 cells in the presence of 5-Ph-IAA, APCin, or a combination of the drugs. **Figure 5D-E** and **S4C-D** show that APCin alone had no statistically significant effect on cell proliferation, in comparison with not treated cells. In contrast, the simultaneous treatment with APCin and 5-Ph-IAA caused a significant reduction in cell proliferation, in comparison with either 5-Ph-IAA or APCin alone.

In summary, our data suggest that NCL abundance directly correlates with the levels of components of the chromosomal segregation machinery in TNBC patient samples and in our in vitro cellular model. Therefore, NCL abrogation has an enhanced anti-proliferative effect in combination with APC/C^Cdc20^ inhibitors, at least in part by impairing DNA dynamics.

## Discussion

Our understanding of nucleolar structural and biological properties has been propelled by the advancement of experimental tools in molecular and cellular biology. However, technical challenges, related to the experimental modulation of their expression, still prevent the molecular characterization of the biological functions of nucleolar proteins.

NCL is one of the most abundant nucleolar proteins, and it has recently gained interest due its altered expression in neurological diseases and cancer^23,28,94–96^. In the present study, we provide *in silico* evidence that NCL overexpression occurs in a broad spectrum of breast tumors, with a significant upregulation in the most aggressive forms of BC, such as TNBC, both at the RNA and protein level (**Figure 1 and S1**). However, this evidence does not exclude the possibility that NCL overexpression can occur also in other cells of the tumor microenvironment, as previously shown^36^.

Like other nucleolar proteins, NCL biological functions remain poorly characterized, due to the lethal effect associated with its chronic downregulation. This is evident from the analysis of the Cancer Dependency Map database not only for BC cell lines, but throughout all the listed cellular models (**Figure S2**). To overcome this limitation and achieve the complete and acute downregulation of endogenous NCL in TNBC cells, we implemented a recently developed technology, the Auxin Inducible Degron (AID) system, in combination with CRISPR/Cas9 knock-in strategies (**Figure 2 and S2**). We observed that endogenous NCL degradation upon OsTIR1-mediated 5-Ph-IAA induced proteolysis can be achieved in approximately 6 h, which is faster than the reported kinetic degradation observed with si/shRNA (3-5 days)^41^. Therefore, our system allows the study of endogenous NCL biological functions with unprecedented temporal resolution, before detectable adaptation or cell death can occur. This degradation kinetic is still slower than previously reported targets of the AID system (30-60 minutes)^43,45,47,54^. As a possible explanation for this slower dynamic, we observed the accumulation of the activated enzyme in the nucleoli monitored as OsTIR1-mEmerald only upon 5-Ph-IAA treatment. This observation suggests that OsTIR-1 mediated NCL ubiquitylation occurs specifically in the nucleolus. The nucleolus is a membrane-less cellular compartment, but physically separated from the nucleoplasm because of the biophysical properties, such as Liquid-Liquid Phase Separation (reviewed in^2^). Therefore, it is conceivable that the reduced accessibility of NCL to OsTIR1 could slow the pace of degradation in comparison with previous studies for non-nucleolar proteins. On the other hand, our data indicate that the AID system could still be functional against phase-separated condensates, such as the nucleolus, which was never studied before.

We assessed the impact of NCL acute abrogation on cellular RNA abundance (**Figure 3C**). Analysis of RNA-sequencing data identified only a small number of genes significantly de-regulated at 16 h upon NCL degradation. This result is not unexpected, as the short time point used for these experiments was specifically selected to identify RNA targets immediately associated with NCL abrogation, and not the result of long-term consequences. Similar findings were also previously reported upon the AID-mediated acute degradation of major regulators of chromatin structure^97^, which could be attributed to the different half-life of mRNA transcripts present in the analyzed cells. Further studies are warranted for a mid- and long-term elucidation of the dynamic effects of NCL abrogation on RNA biology and metabolism, including splicing, capping, poly-adenylation, and translation.

Nonetheless, we observed a specific negative enrichment of genes involved in cell cycle progression, and specifically G2/M transition (**Figure 3C-E**), resulting in impaired cell proliferation upon NCL abrogation. Previous studies, using RNAi-based NCL silencing systems, have reported contradictory results about the role of NCL in the regulation of the cell cycle^41,83,85^.

Conversely, our cell cycle analyses showed that NCL acute depletion causes a marked increase in cells with a tetraploid content of DNA (4N), but a decrease in phospho-Histone H3+ cells suggesting that this phenotype was not due to an increase in mitotic cells (**Figure 4A-D**). Moreover, growth curve analyses showed that the effect on cell proliferation was detectable only after 48 and 72 h from NCL abrogation (**Figure 3A and S3**). We speculated that the discrepancy between NCL degradation and the impact on cell proliferation could be due to a previously unreported role for this protein in regulating the abundance of proteins involved in cell cycle progression at the latest stages of cell division, rather than to a direct positive regulation of the cell division cycle entry. In line with this hypothesis, we found that NCL degradation results in a significant accumulation of bi-nucleated tetraploid cells containing remnants (actin patches) of failed cytokinetic processes (**Figure 4E-F**). These observations also suggest that the increase in cells with a 4N DNA content is not due to their accumulation in the G2 phase of the cell cycle, but rather in a tetraploid bi-nucleated G1 phase.

It has been previously reported that tetraploid cells can attempt further cell division cycles, but the presence of multiple centrosomes leads to the formation of altered mitotic spindles^98–100^. Alterations of centrosome numbers and defective mitotic spindle assembly upon NCL silencing have been previously shown^41^. Therefore, we posit that the observed cytokinesis defects identified in the present study could represent the initial stage of the altered cellular phenotype reported earlier and indicate that the acute degradation mediated by our AID system allows a better temporal characterization of NCL biological functions.

The *in silico* analyses of the TCGA and CPTAC databases showed that NCL expression positively correlates with genes involved in chromosomal segregation, one of the latest crucial steps of the cell division cycle in BC patients (**Figure 5A-B**). Consistently, we showed a significant reduction of proteins regulating chromosomal separation and cohesion in our *in vitro* model, as soon as 24 h after NCL degradation. In addition, NCL-depleted cells displayed a significant increase in sensitivity to inhibitors of the Anaphase Promoting Complex/Cdc20 such as APCin, as expected based on previous reports^92^. In fact, these compounds have previously shown their synthetic lethality with defects of chromatid separation or cohesion^92^.

All in all, our findings support a role for NCL in regulating proteins that regulate cell cycle progression, chromosomal dynamics and segregation. It is therefore tempting to speculate that genetically unstable cancer cells could benefit from increased levels of NCL to prevent mitotic catastrophes resulting in cell death. Future studies will be required to validate this hypothesis and further characterize the mechanisms through which NCL regulates these biological processes in the context of intra-tumoral evolution of aggressive tumors resistant to genotoxic insults. and further characterize the mechanisms through which NCL regulates these biological processes. Moreover, the use of a TNBC cell line displaying dominant-negative mutations of *TP53*^101^ warrants the need of additional validation in multiple cell lines, including normal-like cells. Finally, a dysfunctional cytokinesis could represent one of multiple cellular phenotypes associated with altered NCL expression, as suggested by our GSEA analyses.

In summary, the present study reports the implementation and validation of a novel tool for the acute degradation of nucleolar proteins, which can provide a valuable resource for the study of their biological functions. This system could be used to investigate broader questions about the biology of NCL and of the nucleolus. For example, it can be used to study changes in biophysical properties depending on NCL abundance, or to assess the impact on the biogenesis of rRNA and ribosomes, or to measure the stability of proteins and mRNAs involved in cellular homeostasis during different phases of the cell cycle. Finally, the knowledge gained by this study can inform the development of new therapeutic strategies for TNBC, and potentially for a wide range of human tumors where NCL is over-expressed.

## Materials and Methods

### Plasmids

All the newly generated plasmids reported in this study were created by Gibson Assembly using NEBuilder HiFi DNA Assembly Master Mix (New England Biolabs, E2621L) as per manufacturer’s instructions. The ROLECCS-V2-AS plasmid was previously described^82^. The OsTIR1(F74G)-IRES-H1-mMaroon was generated by replacing the puromycin cassette of Addgene plasmid 140536 (^47^) with the IRES-H1-mMaroon cassette of Addgene plasmid 83842 (pLL3.7m-mTurquoise2-SLBP(18-126)-IRES-H1-mMaroon1)^81^.

To generate the donor plasmids for endogenous *NCL* editing, the genomic region (∼2000 bp) surrounding the natural start codon on exon 1 was cloned into the pUC19 vector (New England Biolabs, N3041S) by Gibson Assembly. Genomic DNA from MDA-MB-231 cells was used as a template to amplify the *NCL* genomic region (Chromosome 2: 231,453,531-231,464,484, Transcript: NCL-201 ENST00000322723.9) of 1 kb upstream and 1 kb downstream the NCL translation start codon. These regions were further used as homology arms for HDR-mediated CRISPR/Cas9-mediated knock-ins. The mAID-mCherry cassette was amplified from Addgene plasmid 72830 (pMK292)^45^ and mAID-HaloTag was amplified from Addgene plasmid 112852^102^. Schematic overview of the tagging vectors is reported in Figure 2A.

To construct the CRISPR/Cas9 NCL-5’ gene targeting vector, a single guide RNA (sgRNA) (5’-CTTCGCGAGCTTCACCATGA-3’) was designed (http://crispr.mit.edu) to target NCL translation start site and the targeting sequence was cloned into pX330-U6-Chimeric_BB-CBh-hSpCas9-hGem (1/110) (Addgene#71707)^103^ according to standard protocols^104^. The same protocol was followed to clone the sgRNA used for *OsTIR1* insertion in the *AAVS1* locus^45,79^.

All the plasmids generated in the study will be available on Addgene (www.addgene.org).

### Cell culture, transfections, and gene editing

MDA-MB-231 cell line was purchased from the American Type Culture Collection (ATCC HTB-26) and cultured in RPMI-1640 medium (Millipore Sigma) supplemented with 10% FBS (Millipore Sigma). Cells were grown in a humidified 37 °C incubator with 5% CO_2_. Identity of cell lines was validated by STR profiling. Cells were regularly tested for Mycoplasma contamination using MycoStripTM Mycoplasma Detection Kit (InvivoGen, #rep-mysnc-10).

To generate NCL-edited cells, 3×10^5^ cells were plated in a six-well plate transfected with the indicated gRNAs and donor plasmids (described above) using Lipofectamine^TM^ 3000 Transfection Reagent (Thermo Fisher Scientific, #L3000001) in Opti-MEM^TM^ I Reduced Serum Medium. (Thermo Fisher Scientific, #31985070). Cells were then grown to a subconfluent T175 cm^2^ flask in medium supplemented with 10% FBS. After incubation with 646-Janelia Fluor® HaloTag® Ligand (Promega, #GA1120), cells were sorted by FACSAria III for both mCherry positive fluorescence at 587 nm excitation, and HaloTag® far red 646 nm positive fluorescence. MDA-MB-231 parental cells and non-stained MDA-MB-231 cells were used as negative controls for mCherry and 646 fluorescence to design the gates. Double positive cells were collected in a 12-well plate and grown as a bulk population until sub-confluence in a T175 cm^2^ was reached. This bulk double-positive population underwent another round of sorting by FACSAria III to collect single cells in 96-well plates. Cells were then screened by fluorescence and western blot analysis for the selection of clones with homozygous expression of AID-NCL edited protein.

To generate NCL-edited cell lines constitutively expressing OsTIR1(F74G), 3×10^5^ AID-NCL edited cells (clone C10) were plated in a 6-well plate and transfected 24 later with donor plasmids ROLECCS V2 AS or OsTIR1(F74G)-IRES-H1-mMaroon, in combination with a plasmid expressing *AAVS1* safe harbor-specific gRNAs. Isolation and validation of single clones were performed as described above. A schematic hierarchy of the established cells lines used in the study is reported in Figure S2F.

### Treatments

To induce the degradation of AID-NCL, cells were treated with 5-phenyl-indole-3-acetic acid (phenyl-auxin, 5-Ph-IAA) (Tocris, #7392). To block the interaction between APC/C and cdc20, cells were treated with 75 μM APCin (2-(2-Methyl-5-nitroimidazol-1-yl)ethyl N-[2,2,2-trichloro-1-(pyrimidin-2-ylamino)ethyl]carbamate, 3-(2-Methyl-5-nitro-imidazol-1-yl)-N-(2,2,2-trichloro-1-phenylamino-ethyl)-propionamide) (Sigma-Aldrich SML1503). Drugs were diluted in culture media at treatment.

### Protein Extraction, Western Blots

For total protein extractions, cells were collected and washed with PBS before adding adequate amount of lysis buffer (1% NP-40, 1 mM EDTA pH 8.00, 50 mM Tris-HCl pH 7.5, 150 mM NaCl) containing a protease and phosphatase inhibitor cocktail (cOmplete^TM^, EDTA-free Protease Inhibitor Cocktail, Millipore Sigma, #COEDTAF-RO; PhosSTOPTM inhibitor tablets, Millipore Sigma, #PHOSS-RO).

Protein concentration was estimated by Bradford assay (Biorad, #5000006). After denaturation at 100 °C for 10 minutes, equal amounts of protein (10-30 μg) were separated using SDS-PAGE, loading samples on 4-20% Mini-PROTEAN® TGX™ Precast Protein Gels (Biorad, #456109). Proteins were then transferred to a 0.45 μm nitrocellulose membrane (Biorad, #1620094) and blocked in 5% non-fat milk or bovine serum albumin in TBST for 1 hour at room temperature. Following blocking, membranes were probed overnight with primary antibodies at 4 °C. The next day, membranes were washed three times with TBST before incubation with secondary antibody at room temperature for 1hour. Detection was performed using Pierce™ ECL Western Blotting Substrate (ThermoFisher Scientific, #32209) and either X-ray blue films, in dark room, or the Li-COR Odyssey FC Imager (Li-Cor Bioscience). When necessary, membranes were stripped using Western BLoT Stripping Buffer (Takara Bio, #T7135A) and re-probed with the desired antibody. Primary antibodies used were anti-NCL (D4C7O) Rabbit mAb (Cell Signaling Technology, #14574); anti-GFP (B-2) Mouse mAb (Santa Cruz Biotechnology, sc-9996); anti-OsTIR1 Rabbit polyclonal Antibody (MBL Bio, #PD048); anti-GAPDH (14C10) Rabbit mAb (HRP Conjugate (Cell Signaling Technology, #12231). Secondary antibodies used were WesternSure® Goat anti-Rabbit HRP Secondary Antibody (Li-Cor Bioscience, # 926-80011), WesternSure® Goat anti-Mouse HRP Secondary Antibody (Li-Cor Bioscience, 926-80010), HRP-conjugated anti-mouse IgG (Millipore Sigma, #NA931V), HRP-conjugated anti-rabbit IgG (Millipore Sigma, #NA934V).

### Incucyte Analyses

For live cell imaging experiments, 2,000-3,000 cells were plated in 96-well plates and time-lapse analyses were performed using IncuCyte S3 Live-Cell Analysis System (Essen BioScience). Images were acquired in a single plane of focus with a 10X objective using bright-field, GFP, and RFP channels, at the indicated timepoints. Each experiment was performed in at least 3 biological replicates. Of each replicate, at least 4 technical replicates were acquired. Quantification of cell confluence and fluorescence integrated intensity was generated by the IncuCyte software. Incucyte raw data were exported as averages of the technical replicates and imported into GraphPad Prism software for statistical analyses.

### RNA sequencing

For RNA sequencing experiment, two independent MDA-MB-231 AID-NCL/OsTIR1 clones were grown in a T175cm flask and either not treated or treated with 2 μM 5-Ph-IAA for 16 hours. Cells were then collected, and RNA was extracted using TRIzol RNA Isolation Reagent (ThermoFisher, #15596026). Library preparation and Next Generation Sequencing were performed at UCLA Technology Center for Genomics and Bioinformatics (TCGB). The library type was Illumina-RNA (rRNA depleted) and Next-Generation Sequencing was performed using Illumina NovaSeq-S1-PE 150 Cycle, with read lengths of 2×150-bp (paired ends). After trimming the library adapter sequences from the raw reads using TrimGalore (github.com/FelixKrueger/TrimGalore), hisat2 (github.com/DaehwanKimLab/hisat2) was used to map the reads to the reference human genome (GRCh38). The output SAM files were converted to BAM files, sorted by index. Reads quality across samples was tested using MultiQC^105^. Gencode gtf file (version 39) was used for gene annotation and counts generation using feature-counts^106^. Raw counts were filtered for low expressed genes following the library size normalization (TMM-Trimmed Mean of M values) using edgeR^107^. The identification of differentially expressed genes between cases and controls was conducted using the R package limma^108^. Adjusted p-values were calculated using the Benjamini and Hochberg method. Heatmaps were generated using the R package pheatmap (https://github.com/raivokolde/pheatmap).

### Data analysis of TCGA, FUSCC, and CPTAC

For TCGA analysis, data was downloaded from The Cancer Genome Atlas (TCGA) database using the Xena browser^109^, provided by University of California Santa Cruz (UCSC). Data from the TCGA Breast Cancer (BRCA) was selected, and samples with PAM50 phenotype designations^66^ were identified. RNA expression data of NCL was obtained for each patient sample. Data points were downloaded, and GraphPad was used to compare subgroups of NCL gene expression and the PAM50 subgroups in order to visualize NCL gene expression levels in the 5 different phenotypes: normal-like, luminal A, luminal B, HER2-enriched, and basal-like. Statistical significance was calculated using Kruskal-Wallis one-way ANOVA with multiple comparisons.

Analysis of NCL expression in different stages of tumor development were analyzed using the same database. Patient samples were categorized based on pathological stage and NCL expression data was obtained for each patient sample. Data was plotted for stages I-IV. Statistical significance was calculated Kruskal-Wallis one-way ANOVA with multiple comparisons.

Similarly, RNAseq FPKM (Fragments Per Kilobase per Million mapped fragments) count data from the FUSCC TNBC cohort were obtained from node OEZ000398 on www.biosino.org.

RNAseq Data for NCL were categorized as basal or non-basal and statistical significance was calculated via two-way Mann-Whitney test.

Protein abundance data for breast cancer patient samples from CPTAC was obtained from cBioportal^110–112^. Protein samples were categorized into PAM50 subgroups and all non-basal tumors were grouped together. Statistical significance was calculated via one-way ANOVA or Mann-Whitney test.

### GSEA

Gene Set Enrichment Analysis (GSEA) was performed on the RNAseq based on normalized count files obtained from the RNAseq comparing samples which were treated and not treated with 2 μM 5-Ph-IAA for 16 hours. Enrichment analysis was performed using the Hallmark Gene Set available from the GSEA database. 1000 permutations were run for each enrichment analysis and statistics were calculated within the GSEA software based on the probability of the obtained calculated enrichment score as compared to the generated permutations. Statistical testing was adjusted for multiple hypothesis testing. For pre-ranked GSEA analysis utilizing patient data, Spearman correlations rho-values between NCL RNA or protein levels and other factors RNA or protein levels were calculated on cBioPortal^110–112^. Genes were ranked from most positively correlated with NCL to most negatively correlated with NCL and this ranked list served as the input to the GSEA. Enrichment analysis was performed using the Gene Ontology Biological Processes Gene Set available from the GSEA database. Enrichment analysis was performed using 1000 permutations as stated above, comparing the generated permutations enrichment scores to that of the input pre-ranked list. Statistical analysis was performed within the GSEA software and statistical testing was adjusted for multiple hypothesis testing.

### Flow Cytometry analyses

MDA-MB-231 AID-NCL/OsTIR1/H1.0-mMaroon1 cells were collected, washed in 1X PBS before being fixed in ice-cold 70% ethanol, and stored at −20 °C overnight. Cells were then pelleted and washed in 1X PBS and incubated with Phospho-Histone H3 (Ser10) (D2C8) XP Rabbit mAb (Cell Signaling Technology, #3377) in 1% BSA/PBS for 90 minutes at room temperature (RT). After primary antibody staining, cells were washed with 1x PBS, and incubated with Goat anti-rabbit Alexa 488 secondary antibody (Life Technologies, #A-11008) in 1% BSA/PBS. Finally, cells were pelleted, washed with 1X PBS, and stained with propidium iodide staining solution (1X PBS containing 10 mg/mL propidium iodide (Thermo Fisher Scientific, #P1304MP), 0.05% Triton X-100 (Millipore Sigma, #9036-19-5), 2.5 µg/mL RNAse A (Thermo Fisher Scientific, # EN0531)). Flow cytometry analysis was performed using LSRFortessa (Beckman-Dickinson) Flow Cytometer, modeling at least 5,000 events per sample. Gating was designed to quantitate the absolute population of cells with a 4N content of DNA, and either positively (4N; pH3+) or negatively (4N; pH3-) stained for pH3(S10) antibody, using the FloJo software. Cell cycle analyses (based on PI staining alone) were performed with ModFit software, v5.0. Two independent clones were analyzed, for a total of three experiments. Each experiment was performed in technical duplicate, and data were normalized for the untreated controls. Significance was calculated using GraphPad Prism software. Welch’s t-test was applied.

### Immunofluorescence

The formation of bi-nucleated cells was evaluated by immunofluorescence as follows. Cells were plated on sterile glass coverslips (Gold Seal Cover Glass, thickness 1.5, Thermo Fisher, #3406) pretreated with poly-L-ornithine solution (Sigma Aldrich, #P4957) in 6-well plates in regular medium. The next day, cells were treated with 2 µM 5-Ph-IAA for 72 hours or left untreated. Following treatment, medium was aspirated, and coverslips were washed with 1X PBS and fixed with cold 4% PFA/PBS pH 7.4 (Paraformaldehyde Solution, 4% in PBS, Thermo Scientific™, #J19943.K2) for 15 minutes at RT. Fixative solution was then removed, and cells were incubated with 0.3 M Glycine solution (Glycine USP, Gojiara Fine Chemicals, #GC1004, dissolved in dH_2_O and filtered) for 5 minutes at RT. Coverslips were then rinsed with 1X PBS and incubated with 1X PBS for 5 minutes at RT. Cells were then permeabilized with 0.2% Triton X-100 in 1X PBS for 10 minutes at RT. Following a wash with 1X PBS, cells were blocked in 5% BSA/PBS (filtered) for 1 hour at RT. Next, coverslips were incubated with anti-Lamin A/C (4C11) Mouse mAb (Cell Signaling Technology, #4777), diluted in 1% BSA/PBS filtered solution, for 1 hour at RT in a covered humidified chamber. Coverslips were then rinsed with 1X PBS and washed two times for 10 minutes each with 1X PBS. Next, cells were incubated with Goat anti-Mouse IgG Antibody AlexaFluor™ 488 (Fisher Scientific, #A11029) and Phalloidin-iFluor 647 (Abcam, #ab176759) in 1% BSA/PBS filtered solution for 1 hour at RT in a covered humidified chamber. Coverslips were then rinsed with 1X PBS and washed three times for 10 minutes each with 1X PBS. Coverslips were mounted on Superfrost Plus Microscope Slides (Fisherbrand, #12-550-15), using VECTASHIELD Antifade Mounting Medium with DAPI (Vector Laboratories, #H-1200-10). Slides were left at RT overnight, before evaluation. Images were captured in a single plane of focus at 60x magnification using an EVOS™ M5000 Imaging System (EVOS™ M5000 Software). Bi-nucleated cells were defined as single cells identifiable by an actin border positive to phalloidin staining (magenta), with two separate nuclei encased by nuclear envelope positive to Lamin A/C staining (green). For quantification purposes, at least 15 independent fields were captured, for a total of at least 100 cells for each sample. The percentage of bi-nucleated cells per image was used to generate the comparison between treated and not treated cells. Data were analyzed using GraphPad Prism software. Statistical significance was calculated using one-way ANOVA with Sidak’s multiple comparison test.

### TMT quantitative proteomic analyses

Quantitative proteomics was performed as previously described^113,114^. Briefly, cells were plated in T175cm flasks and grown to subconfluency. Cells were either subjected to 2 µM 5-Ph-IAA for 24 hours or remained untreated. Cell lysis was performed using 8M Urea, 50 mM Triethylamonium bicarbonate (TEAB) buffer containing protease and phosphatase inhibitors (Roche), as described in “Protein Extraction”. The samples were then sonicated for 30 seconds on ice (amplitude 25%), and concentration was estimated Pierce^TM^ BCA Protein Assay Kit (Thermo Fisher Scientific, Cat. 23225). Tryptic digestion, Tandem Mass Tag (TMT) labelling, fractionation, mass spectrometry, and validation analyses were performed by Bioinformatics Solutions Inc., using at least 100 μg of extracted proteins. Peptide spectra were searched using the Uniport validated human proteome and Peaks Studio (v11), using a 15-ppm precursor ion tolerance for peptide mapping. A decoy fusion method was employed to determine if peptide-spectrum matchings are true matchings or likely false positives. A cut-off of 1% false-discovery-rate was used to filter poor peptide matchings. Relative abundance (Log2 Fold Change) and significance (−10Log10 of *p*-value) were calculated by comparing the peptide quantification values control and 5-Ph-IAA treated samples, via Peaks Studio. Data were considered significant for −10Log10 of *p*-value >13.

### Statistical analyses

The number of replicates, and the statistical tests that were used for each experiment are specified in the relative figure legend. For statistical analysis, GraphPad Prism software was used. Data was considered statistically significant for *p* < 0.05.

## Supplementary Data

A detailed description of supplementary figures is reported in the Supplementary Information file.

## Data availability

RNA-sequencing and TMT proteomics data are pending deposition into public data repository. Source data files are available in the Supplementary Data or from the Principal Investigator upon request.

## Acknowledgements

This work was supported by seed funds from The Ohio State University Comprehensive Cancer Center (DP), the President’s Research Excellence Accelerator Award (DP, AED, HS) and ORIP-NIH (K01OD031811-01) (AED). The Gene Editing Shared Resource, the Flow Cytometry Shared Resource and the Genomics Shared Resource that contributed to this study are supported by the Cancer Center Support Grant P30CA016058. We also thank the countless investigators who made their plasmids and reagents available through public repositories such as Addgene. The results shown here are in part based upon data generated by the TCGA Research Network (https://www.cancer.gov/tcga) and the CPTAC Clinical Proteomic Tumor Analysis Consortium (https://proteomics.cancer.gov/programs/cptac). Figures were partially generated using BioRender (www.biorender.com).

## Author Contributions

DP, AT, and JM conceived the project, designed, performed, experiments, analyzed data, wrote the manuscript, and prepared the figures. AT and JM contributed equally; therefore they can list this manuscript in their records swapping their names. VA performed analysis of proteomics data, provided extensive support in experiment design and in the analysis of external databases. DSK, NSR, ZZ, SL, and LR provided support for the experiments and for the generation of the cell lines used in this project. IC and ABR performed bioinformatic analyses of RNA-sequencing data. EC, TJM, AED, HS, VC, and CMC provided guidance in project and experiment design and in data collection and interpretation. All the authors have read and approved the final version of the manuscript. AT and JM contributed equally to the project.

## Supplementary Data

### Supplementary Figures and Supplementary Figure Legends

**Figure S1.**
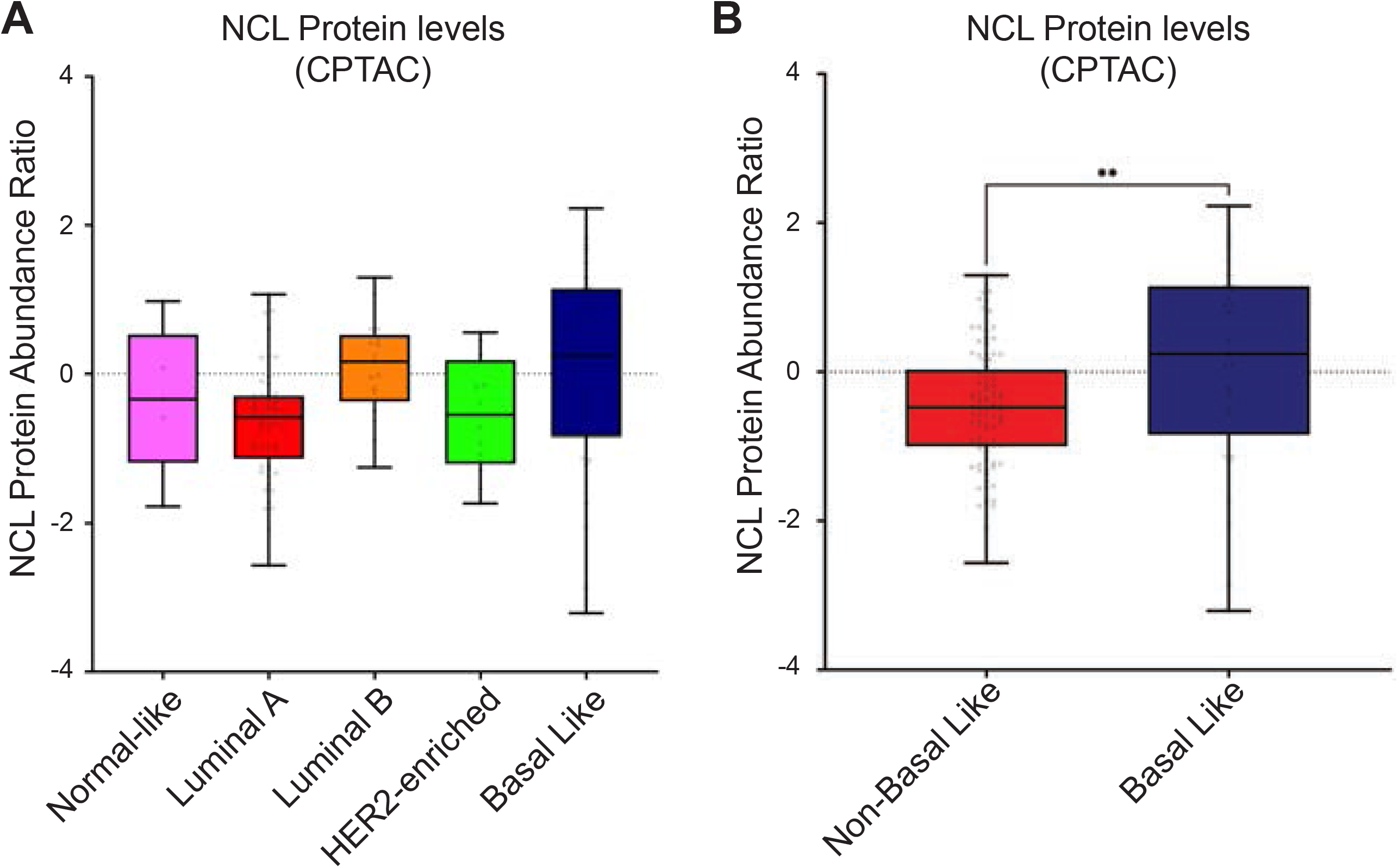
Increased NCL protein levels in Basal Like breast tumors. **A)** NCL protein levels among different breast tumor subtypes compared to the normal-like breast cancer samples. Protein levels were obtained from 122 patient samples (Normal-like, n=5; Luminal A, n=57; Luminal B, n=17; HER2-enriched, n=14; Basal Like, n=29 available through the Clinical Proteomic Tumor Analysis Consortium (CPTAC). **B)** Comparison of NCL protein levels in Basal Like (n=29) vs pooled non-Basal Like (n=93) BC, as shown in A. Significance was defined using Mann-Whitney test. **: *p* < 0.01.

**Figure S2.**
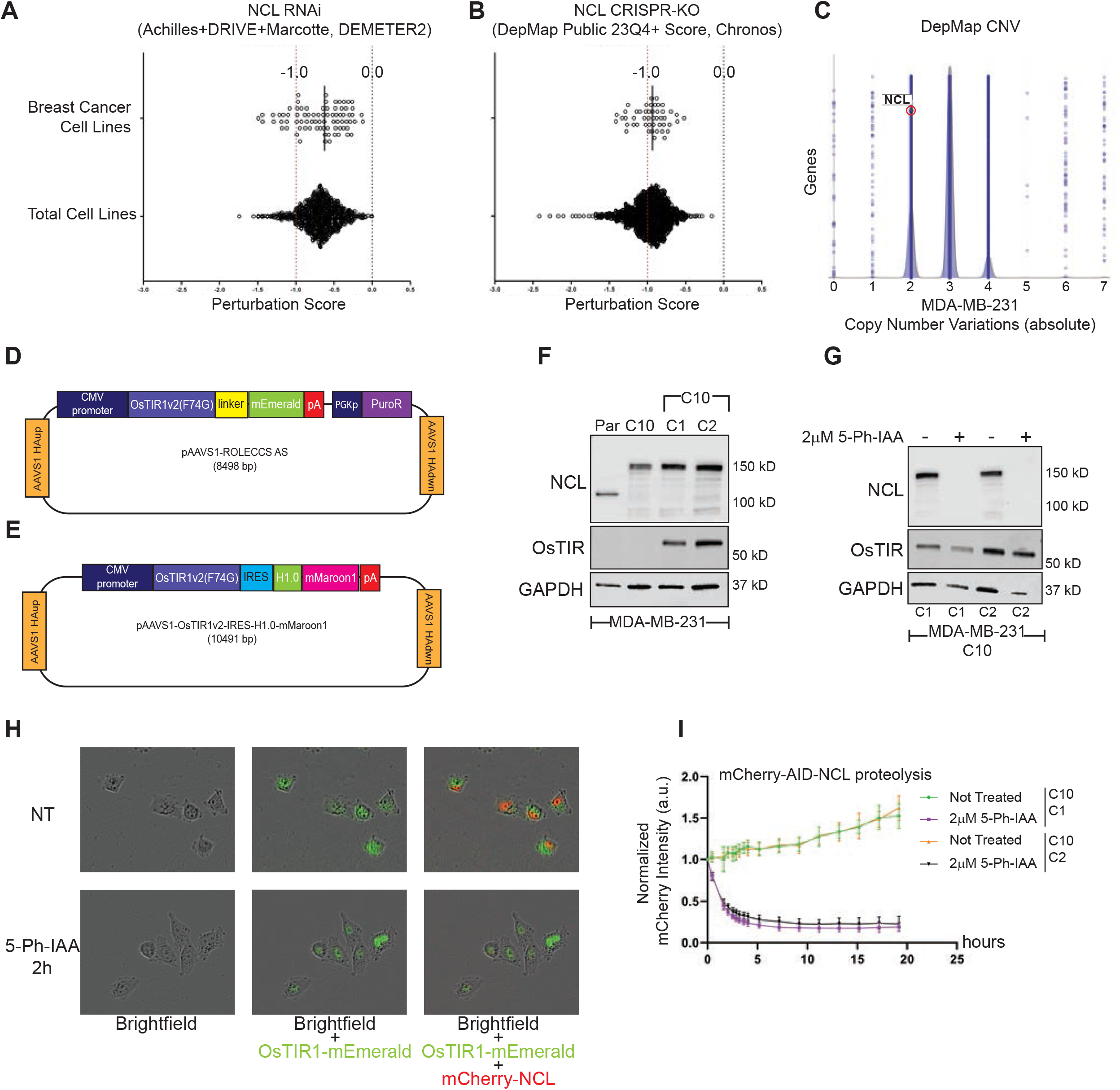
Development of an Auxin Inducible Degron for NCL. **A-B)** Cancer DepMap data from RNAi (**A**) and CRISPR-KO (**B**) screen databases (as indicated) show NCL Perturbation Score across Breast Cancer cell lines (empty dots) or total cell lines (black dots). **C)** Copy number variation (CNV) analysis of genes in MDA-MB-231 cells from the Cancer DepMap data. Blue dots represent individual genes, aligned based on their absolute copy number. Nucleolin (*NCL*) is highlighted in red. **D)** Schematic representation of the plasmid used as a donor for *AAVS1* site-specific integration of the *OsTIR1*(F74G)-mEmerald fusion gene (used for the generation of clones C10.C4 and C10.D9). **E**) Schematic representation of the plasmid used as a donor for *AAVS1* site-specific integration of the vector co-expressing *OsTIR1*(F74G) and histone H1-mMaroon1 fusion protein (used for the generation of clones C10.C1 and C10.C2). IRES: internal ribosome entry site. **F-G)** Western blot analyses for effective integration (F) and biological activity (G) of OsTIR1(F74G) in MDA-MB-231 AID-NCL homozygous clones (see also Figure 2 and S2E) using the indicated antibodies. GAPDH was used as loading control. For degradation experiments, clones were treated for 24 h with 5-Ph-IAA. **H)** Representative images of Incucyte experiments on MDA-MB-231 mCherry2/Halo-AID-NCL/OsTIR1-mEmerald cells (see also Figure 2G). The RFP channel was used to detect mCherry2-AID-NCL, and the GFP channel was used to assess OsTIR1-mEmerald expression and localization. Cells were left untreated or treated with for 2 h with 5-Ph-IAA (see also Figure 2G). **I)** Quantitative analysis of mCherry/NCL intensity from Incucyte experiments on the indicated cell clones, not treated, or treated with 5-Ph-IAA. RFP integrated intensity per image was normalized for t=0. The experiment was performed on 9 biological replicates, with 5 technical replicates. Error bars show standard deviation.

**Figure S3.**
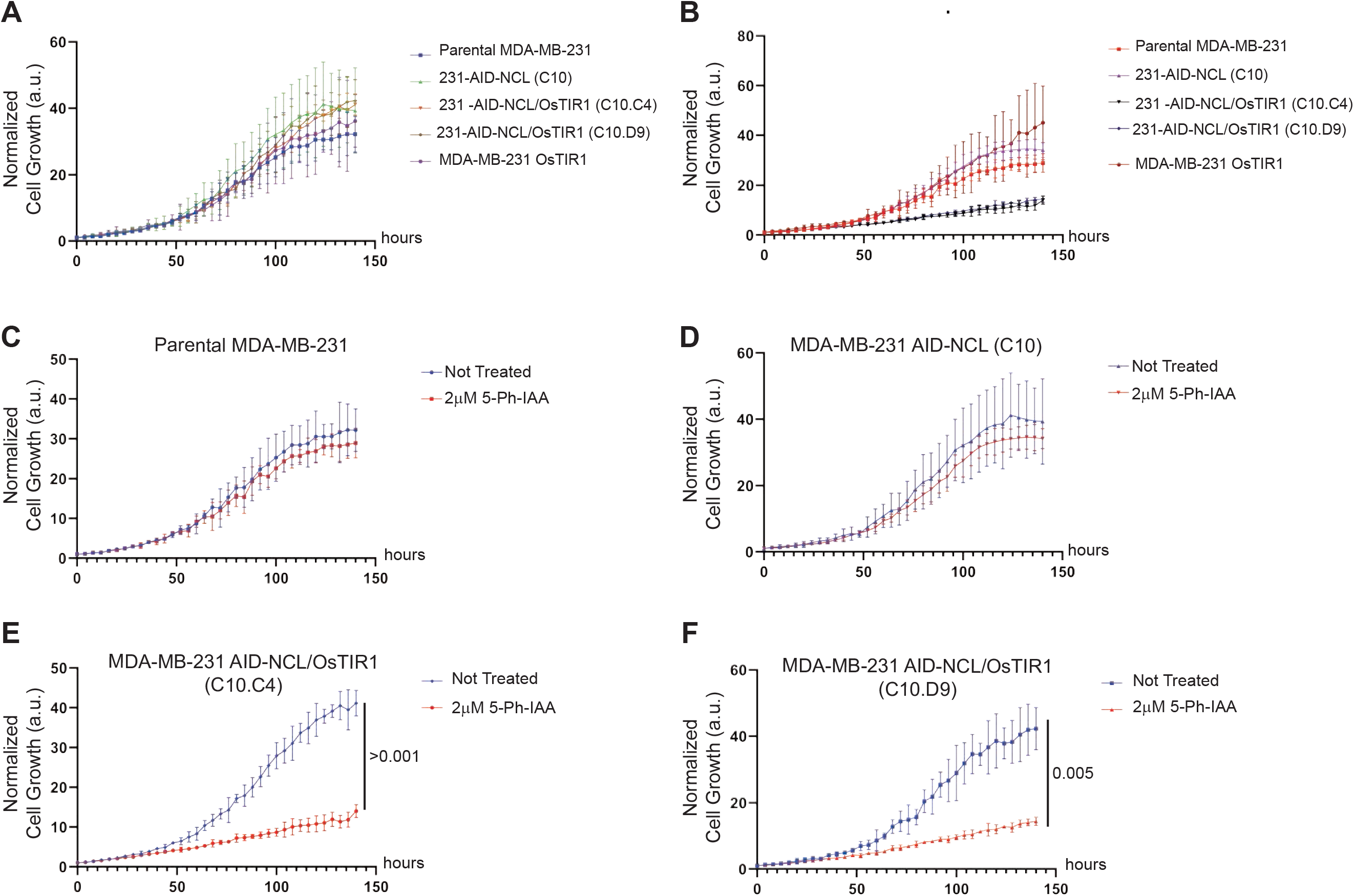
AID-dependent NCL abrogation affects cell proliferation. **A-B)** Growth curves comparing parental MDA-MB-231 and all the gene-edited derivative clones, left untreated (A) or treated with 2 μM 5-Ph-IAA (B). **(C)** Growth curve of parental MDA-MB-231 cells, with or without treatment with 2 μM 5-Ph-IAA, **(D)** Growth curve of MDA-MB-231 AID-NCL clones (not expressing OsTIR1) with or without treatment with 2 μM 5-Ph-IAA. **(E-F)** Growth curve of two AID-mCherry2-NCL OsTIR1 MDA-MB-231 clones, with or without treatment with 2 μM 5-Ph-IAA for up to 140 hours. All the growth curves in this experiment and in main Figure 3 are summarized in Figure S3A-B, and individual comparisons are shown in independent panels. Cell proliferation was monitored with Incucyte live-cell imaging system. All the growth curves were performed in 5 biological replicates with 5 technical replicates. Error bars show standard deviation. Statistical significance was calculated using 2-way ANOVA with Tukey’s multiple comparison test. Where significant, the exact *p*-value is reported (E-F).

**Figure S4.**
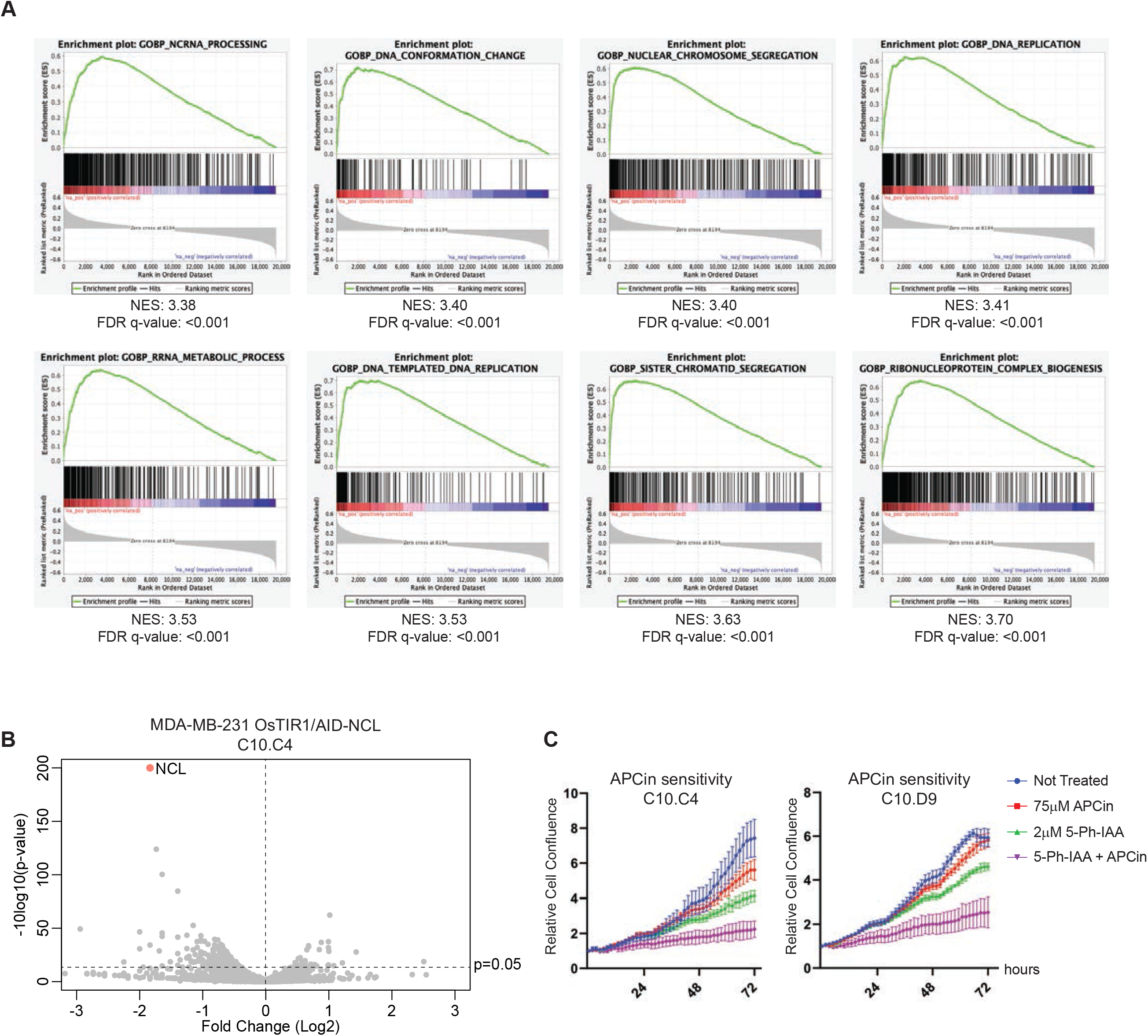
NCL expression correlates with genes involved in RNA metabolism and chromosomal dynamics in BC patients, and its abrogation enhances sensitivity to APC inhibitors. **A)** Enrichment plots of eight among the top ten most enriched GO terms containing genes positively correlating with NCL RNA expression in BC patients (see also Figure 1A and Figure 5A). **B)** Volcano plot of differentially abundant proteins upon NCL abrogation in MDA-MB-231 OsTIR1/AID-NCL cells (clone C10.C4), detected by TMT-proteomic analysis. Reported data is the full range of detected proteins as in Figure 5, with a maximum −10log_10_(p-value) of 200. **C)** Incucyte growth curves, referred to end-point experiments shown in Figure 5D-E, of two independent MDA-MB-231 OsTIR/AID-NCL clones, left untreated or treated with indicated doses 5-Ph-IAA, APCin, or combination, for up to 72 h. Incucyte live-cell imaging system was used to monitor cell proliferation. Cell confluence was normalized for t=36’. The experiment was performed in three biological replicates, with five technical replicates. Error bars show standard deviation.

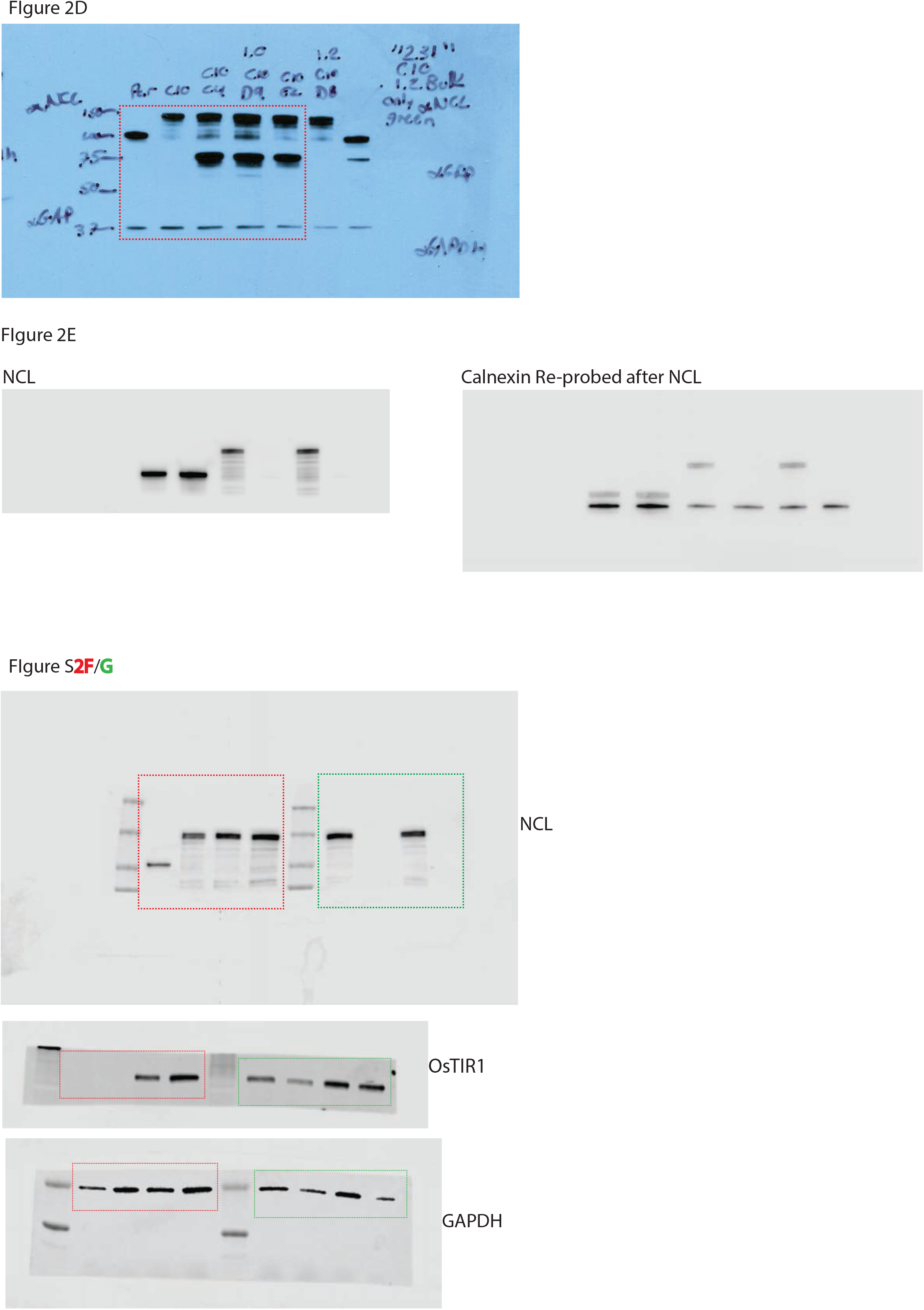

**Table S1:** Deregulated proteins upon NCL acute degradation (GO Term: Ribosome Biogenesis)

| Protein Name | FC (Log2) | Significance - $10\log_{10}(\text{p-value})$ |
| --- | --- | --- |
| RPL14 | -1.4 | 84.7 |
| RPS9 | -1.3 | 43.4 |
| RPS4X | -0.8 | 39.0 |
| RPS16 | -1.0 | 37.9 |
| RPS24 | -0.7 | 34.1 |
| RPS7 | -0.7 | 33.7 |
| RPS13 | -0.8 | 30.6 |
| RPL6 | -0.8 | 30.3 |
| RPL7A | -0.7 | 28.6 |
| RPL10A | -0.6 | 28.1 |
| RPS5 | -0.6 | 28.1 |
| RPS23 | -0.6 | 25.4 |
| RPL7 | -0.8 | 24.6 |
| RPL10 | -0.7 | 22.0 |
| RPS15A | -0.7 | 20.4 |
| RSL1D1 | -0.6 | 19.1 |
| RBM34 | -0.8 | 17.9 |
| EXOSC3 | 0.6 | 17.8 |
| RPS11 | -0.5 | 17.8 |
| SERBP1 | 0.5 | 17.7 |
| RPS6 | -0.6 | 17.6 |
| RPS17 | -0.5 | 16.4 |
| RPS8 | -0.5 | 16.3 |
| RPL35 | -0.7 | 15.1 |
| RPL11 | -0.5 | 14.9 |
| DDX52 | -0.8 | 14.9 |
| UTP3 | 0.8 | 14.8 |
| RPL5 | -0.4 | 14.3 |
| UTP14A | -0.7 | 14.2 |
| NAT10 | -0.4 | 13.8 |
| EXOSC8 | 0.5 | 13.8 |

**Table S2:** Deregulated proteins upon NCL acute degradation (GO Term: Chromosome Segregation)

| Gene | FC (Log2) | Significance - $10\log_{10}(\text{p-value})$ |
| --- | --- | --- |
| SLC25A5 | -1.6 | 45.5 |
| KIF2A | -1.4 | 26.3 |
| MAP1S | -1.2 | 19.6 |
| SMARCD1 | -0.9 | 18.7 |
| TPX2 | -1.2 | 18.2 |
| EML4 | -0.6 | 16.9 |
| RPS3 | -0.4 | 13.7 |

